# Clofazimine targets essential nucleoid associated protein, mycobacterial integration host factor (mIHF), in *Mycobacterium tuberculosis*

**DOI:** 10.1101/192161

**Authors:** Wanliang Shi, Shuo Zhang, Jie Feng, Peng Cui, Wenhong Zhang, Ying Zhang

## Abstract

Clofazimine (CFZ) is a phenazine derivative used for treatment of leprosy, MDR-TB and XDR-TB. There is recent interest in understanding how CFZ works following the demonstration of its unique ability to shorten the treatment of MDR-TB. However, the target of CFZ in mycobacteria has remained elusive. Here, we show that CFZ binds to mycobacterial integration host factor (mIHF), which is an essential nucleoid associated protein in mycobacteria involved in DNA protection, chromosome organization and global gene regulation. We demonstrate that CFZ inhibits mIHF binding to DNA and interferes with mycobacterial gene expression. This mode of action is unique among all antibiotics including antimycibacterial agents and may help to explain its unusual action against *Mycobacterium tuberculosis*. Our study provides new insight about the mechanism of action of this intriguing drug and has implications for developing more effective treatment of TB.

---

Although clofazimine (CFZ) was discovered as an anti-tuberculosis drug in 1957^1^ and later used for treatment of leprosy^2^, multi-drug-resistant tuberculosis (MDR-TB)^3^ and extensively drug-resistant tuberculosis (XDR-TB)^4^, the exact mechanism of action of CFZ is poorly understood^5,6^. Various studies suggest that CFZ seems to have multiple effects such as binding to DNA^7^, interfering with redox cycling^8^, causing membrane destabilization and dysfunction^9^, and production of reactive oxygen species^10^. To identify potential targets of CFZ, we used a proteomic approach to find proteins that could bind to CFZ by affinity chromatography (fig. s1 and fig. s2), a strategy which we previously used to successfully identify a target of pyrazinamide^11^.

Lysates of *M. tuberculosis* were allowed to interact with the CFZ-coupled agarose beads and the control agarose beads in columns by affinity chromatography, followed by analysis of the eluted fractions on SDS-PAGE. Several protein bands eluted from the CFZ-agarose column were seen but no visible band was found from the control column (Fig. 1). Mass sepctrometry analysis of the candidate proteins that were bound to CFZ identified four ribosomal proteins (RplD, RpsP, RpsQ and RplD) and the mycobacterial integration host factor (mIHF, Rv1388). Based on Coomassie Blue staining, mIHF was the most abundant protein with an apparent molecular weight of about 12 kDa (Fig. 1).

**Figure 1.**
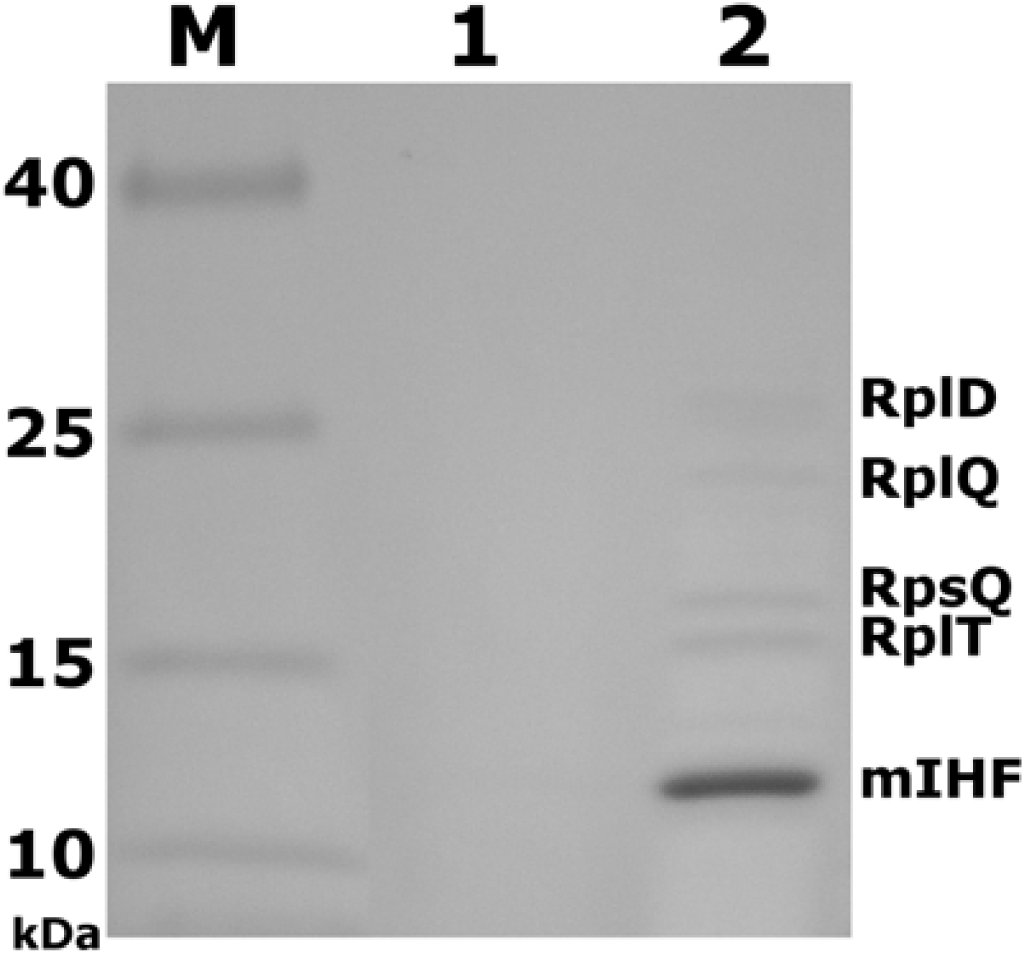
Potential target proteins binding to CFZ analyzed by SDS-PAGE. *M. tuberculosis* whole-cell lysates were loaded onto the CFZ-linked and blank control columns, and the proteins that bound to the control agarose bead column (Lane 1) and CFZ-linked column (Lane 2) were analyzed by SDS-PAGE with Lane M as the protein molecular weight marker. The protein bands were excised and subjected to mass spectrometry analysis, which identified these proteins as RplD, RplQ, RpsQ, RplT and mIHF, respectively.

The annotated mIHF from the *M. tuberculosis* H37Rv genome database indicates that its full length is 190 amino acid residues^12^ (fig. s3). On the other hand, mIHF from *M. tuberculosis* when overexpressed as either a C-terminal 111 aa or 190 aa full length protein was shown to be able to bind DNA and compact DNA in vitro^13,14^. To determine which is the correct size of the mature protein, we cut out the 12 kDa band from the gel (Fig. 1) and subjected it to N-terminal amino acid sequencing by Edman degradation and identified the first 5 amino acid residues to be –ALPQL. This finding indicates that the mIHF from *M. tuberculosis* is in fact 105 aa, rather than the 190 aa as annotated in the database, and is in keeping with the size of the *M.* s*megmatis* mIHF^15^.

The mIHF is rich in strongly basic residues (22/105 residues in lysine and arginine) with a calculated isolectric point 9.91 and its secondary structure was rich in helical content. These features are consistent with the finding that mIHF is a DNA binding protein^13^. To investigate if 105 aa mIHF of *M. tuberculosis* binds to DNA, the recombinant *M. tuberculosis* mIHF was overexpressed in *E. coli* strain BL21 (DE3) and purified (fig. s4). The purified mIHF interacted with both circular and linear plasmid DNA in a concentration dependent manner as shown by electrophoretic mobility shift assay (EMSA) (fig. s5).

To assess whether mIHF interacts with CFZ directly, a gel shift assay was used to analyze the complex formation between the recombinant *M. tuberculosis* mIHF and CFZ. The acidic native gel was used to separate native basic proteins^16^, and the complex formation of native mIHF and CFZ was visualized on acidic native gels. The results showed that the mIHF and CFZ formed a complex (Fig. 2a). However, no complex formation was observed with the control TB drug rifampin (RIF) which has similar color as CFZ (Fig. 2b) or with another control, oil red O, which is a fat-soluble dye with similar characteristics as CFZ (Fig. 2c), even at a high concentration up to 500 μg/ml. Therefore, CFZ interacted with *M. tuberculosis* mIHF specifically.

**Figure 2.**
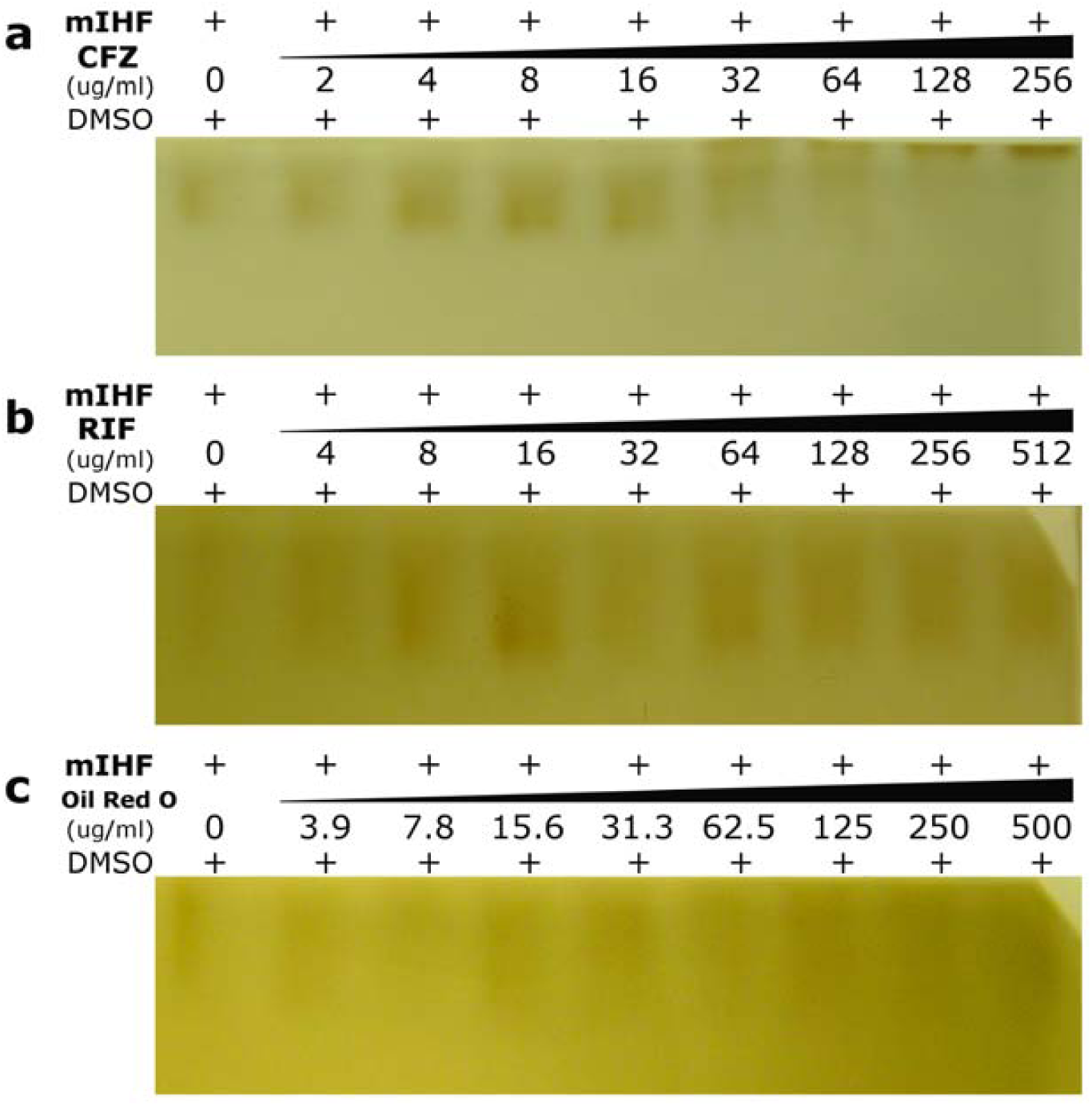
Complex formation between CFZ and mIHF as shown by acidic native PAGE gel. **a**, Concentration-dependent complex formation of CFZ and mIHF. mIHF (180 ng) was incubated with CFZ at different concentrations at 0, 2, 4, 8, 16, 32, 64,128 and 256 μg/ml (from left to right side, repectively) in 15 μl reaction. **b**, Rifampin (RIF) as a control drug did not interact with mIHF. mIHF (180 ng) was incubated with RIF at different concentrations at 0, 4, 8, 16, 32, 64,128, 256 and 512 μg/ml (from left to right side, repectively) in 15 μl reaction. **c**, The control group oil red O did not interact with mIHF. mIHF (180 ng) was incubated with oil red O at different concentrations at 0, 3.9, 7.8, 15.6, 31.3, 62.5,125, 250 and 500 μg/ml (from left to right side, repectively) in 15 μl reaction.

To further examine the effect of CFZ on mIHF binding to DNA, mIHF was incubated with different concentrations of CFZ (Fig. 3a), with RIF (Fig. 3b) or oil red O (Fig. 3c) as controls, respectively. As shown in Fig. 3a, CFZ inhibited mIHF and DNA complex formation in a concentration dependent manner (Fig. 3a, Lane 2-13). In contrast, the control drug RIF (Fig. 3b, Lane 2-13) and oil red O (Fig. 3c, Lane 2-13) had no effect on mIHF-DNA interaction. In addition, CFZ alone did not cause altered DNA migration on agarose gel (fig. s6), indicating CFZ does not bind to DNA directly, but instead binds to DNA indirectly via mIHF. This is in contrast to previous studies that claimed CFZ directly binds to DNA^7,17,18^.

**Figure 3.**
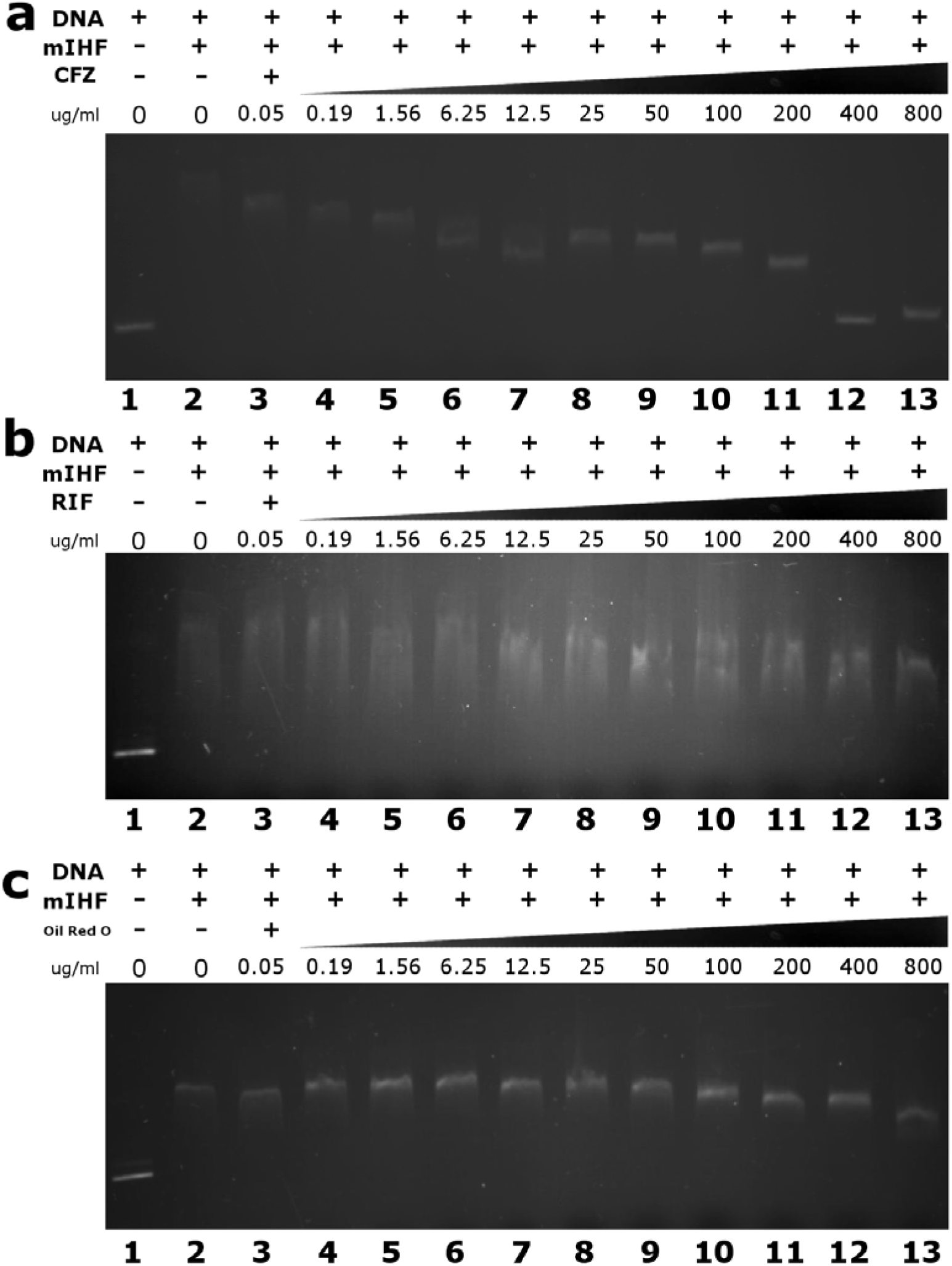
CFZ inhibits interaction of mIHF binding to DNA. **a**, Concentration-dependent inhibition of mIHF binding to linear plasmid DNA by CFZ. Linear pUC19 plasmid DNA (50 ng) was used as the control without mIHF and CFZ (Lane 1). The mIHF (50 ng) was incubated with different concentrations of CFZ from 0 to 800 μg/ml on ice for 15 min, and the mIHF DNA mixture was run on 0.5% agarose gel to visualize the CFZ inhibition of mIHF interaction with DNA stained by EtBr (Lanes 2 to 13). **b**, No inhibition of mIHF binding to linear plasmid DNA by RIF. DNA was used as the control without mIHF and RIF (Lane 1). The mIHF was incubated with different concentrations of RIF from 0 to 800 μg/ml on ice for 15 min and analyzed as above (Lanes 2 to 13). **c**, No inhibition of mIHF binding to linear plasmid DNA by oil red O. DNA was used as the control without mIHF and oil red O (Lane 1). The mIHF was incubated with different concentrations of oil red O from 0 to 800 μg/ml on ice for 15 min and the mixture was analyzed as above (Lanes 2 to 13).

To investigate whether CFZ affects mIHF behavior in vivo in mycobacterial cell, a fusion protein of mIHF with red fluorescent protein (RFP) was constructed under the control by a tetracycline inducible promoter. The mIHF-RFP fusion protein was found to localize at nucleoid zone in *M. tuberculosis* H37Ra/pUV15tet-mIHF-RFP strain at log phase when inducer, anhydrotetracycline (ATc), was added (Fig. 4 a and c). The mIHF-RFP protein was localized at nucleoid zone as a tight spot. However, clusters of red fluorescence were diffused to the whole cell when the mIHF-RFP induced culture was exposed to CFZ (Fig. 4 b), indicating that CFZ changed mIHF location in the *M. tuberculosis* cell. To evaluate whether mIHF location is altered by CFZ, *M. tuberculosis* H37Ra/pUV15tet-mIHF-RFP strain was exposed to different concentrations of CFZ first and then the expression of mIHF-RFP was observed by microscopy upon tetracycline induction. Interestingly, the red fluorescence was diminished depending on the CFZ concentration. There were fewer fluorescent cells and lower fluorescent density in the 1 μg/ml (Fig. 4 d) and 5 μg/ml (Fig. 4 e) CFZ treated groups than the untreated control group (Fig. 4c). Furthermore, no visible red spot at nucleoid zone was seen with the 25 μg/ml CFZ treated group (Fig. 4 f). We found that CFZ did not cause cell death in vitro under this condition, which is consistent with the previous observation^19^.

**Figure 4.**
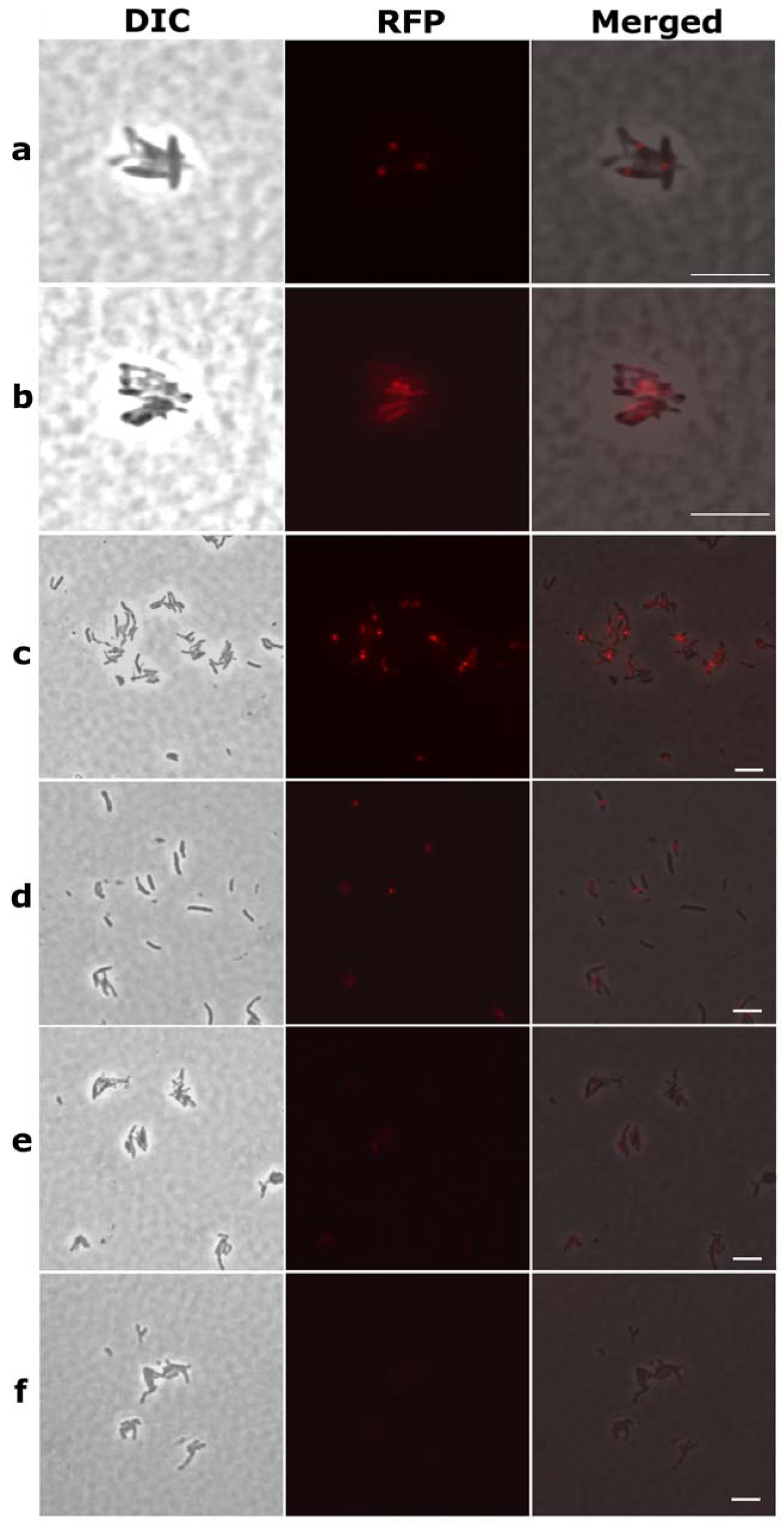
CFZ alters the localization of mIHF in *M. tuberculosis*. **a**, mIHF-RFP fusion protein was induced by ATc (5 ng/ml) for 6 days in *M. tuberculosis* H37Ra/pUV15tet-mIHF-RFP strain. **b**, mIHF-RFP fusion protein was induced by ATc for 3 days and then followed by 5 μg/ml CFZ exposure for 3 days in *M. tuberculosis* H37Ra/pUV15tet-mIHF-RFP strain. *M. tuberculosis* H37Ra/pUV15tet-mIHF-RFP strain was exposed to 0 μg/ml (**c**), 1 μg/ml (**d**), 5 μg/ml (**e**), and 25 μg/ml (**f**) of CFZ for 3 days followed by induction by 5 ng/ml ATc for 3 days. Differential interference contrast (DIC), red fluorescent protein (RFP) and merged images are presented. The scale bar represents 10 μm.

Previously, it was shown that CFZ affected gene expression in *E. coli* and *Staphylococcus aureus*^17^. Furthermore, CFZ inhibited the growth cycle of mycorobacteriophage in *M. aurum*^17^, whose mIHF shares 98% identities with the *M. tuberculosis* mIHF. mIHF is involved in DNA compaction and gene regulation^13^. Although mIHF is an abundant protein in *M. tuberculosis*, severe growth defect occurs when the mIHF level was reduced to lower than 10% while complete depletion of mIHF caused cell death^20^. Thus, CFZ inhibition of mIHF function and its interference with gene expression may cause growth inhibition at low CFZ concentrations but could cause cell death at high CFZ concentrations or longer exposure time. Further studies are needed to determine the molecular and structural basis for this mode of action.

While overexpression of drug targets often causes drug resistance, we found that overexpression of mIHF did not cause significant CFZ resistance in *M. tuberculosis* compared with the parental strain and empty vector control strain. The same phenomenon was observed with the target of linezolid, RplC whose overexpression did not cause resistance to linezolid^21^. In fact, overexpression of the drug target could have a wide range of effects in terms of resistance to that drug, ranging from increase, unchanged, to even decrease ^22^. Since mIHF is an abundant protein that is present at the same high levels in all growth phases in *M. tuberculosis*^20^, any exogenous overexpression on a very high background levels of mIHF would be expected to produce hardly any additional resistance. Further studies are needed to address whether mIHF expression levels contribute to or correlate with CFZ resistance. Although mutations in three genes, Rv0678, Rv2535c and Rv1979c, are typically associated with CFZ resistance^23,24,25^, no mIHF mutation was found in those CFZ resistant mutants. Future studies are necessary to systematically characterize CFZ-resistant clinical strains to determine the role of mIHF gene in mechanism of CFZ resistance in *M. tuberculosis*.

mIHF is a small, heat-stable protein that promotes the integrative recombination of phage DNA into the chromosome through formation of specific integrase-mIHF-attP complexes in mycobacteria There is only one copy of mIHF in mycobacteria which is quite distinct from host integration factor in *E. coli*^15^. mIHF is essential for *M. smegmatis* viability and *M. tuberculosis* growth^26,27^. That mIHF binding to DNA and bending DNA^13^ suggests that it is a global gene expression regulator in *M. tuberculosis* with more functions than that of *E. coli*. Many studies indicated that the interaction between IHF and DNA is complex, with IHF binding to DNA by different modes that induce different DNA bending patterns, and these DNA binding modes are sensitive to different conditions^28,29^. Our findings that mIHF binds to DNA and that CFZ affected mIHF binding to DNA and altered mIHF location in vivo suggest that CFZ could inhibit gene expression by affecting mIHF and DNA interaction in *M. tuberculosis* causing the cell to be more susceptible to secondary attacks by other drugs or reactive oxygen species (ROS) or nucleases that damage DNA. Such a mechanism of action is consistent with recent observation that CFZ activity is enhanced mainly by drugs such as fluoroquinolone, aminoglycoside, and acid pH and PZA that could produce ROS or damage DNA^30^. Further studies are required to test this possibility in the future. Our study provides novel insights about the unique mechanism of action of CFZ and implicates mIHF as a novel target for developing new drugs for overcoming drug resistance and more effective treatment of persistent mycobacterial infections.

## Methods

### Preparation of CFZ affinity chromatography column

CFZ was coupled to the immobilized diaminodipropylamine (DADPA) resin (G-Biosciences, St. Louis, MO, USA) to generate an affinity chromatographic matrix according to the manufacturer's instructions. Briefly, 10 ml of immobilized DADPA resin was centrifuged at 700 g for 2 minutes to remove the storage buffer. The resin was washed with coupling buffer (0.1M MES, 0.15M NaCl, pH4.7) twice. After washing, the resin was mixed thoroughly in the coupling buffer containing 0.2 μg/ml CFZ and 1 ml 37% formaldehyde. The column was incubated with gentle agitation at 37 °C overnight. Then the column was washed to remove excess unbound ligand using coupling buffer. The control column was coupled with 37% formaldehyde without CFZ. The columns were washed with three cycles of alternating pH with 0.1 M acetate buffer (pH 4.0) containing 0.5 M NaCl followed by a wash with 0.1 M Tris-HCl buffer pH 8 containing 0.5 M NaCl.

### Protein lysate preparation

*M. tuberculosis* H37Ra was cultured in Sauton’s medium at 37°C with gentle shaking for three weeks. Sauton’s medium consisted of the following composition (per liter): 4 g of L-asparagine, 0.5 g of monopotassium phosphate, 0.5 g of magnesium sulfate, 50 mg of ferric ammonium citrate, 2 g of citric acid, 1 mg of zinc sulfate, and 60 ml of glycerol (with 0.05% Tween 80 added after sterilization). Mycobacterial cells were collected and washed with PBS buffer (pH 7.0) twice. Mycobacterial protein lysates were prepared by sonication followed by collecting the supernatants after centrifugation as described previously^11^. The protein concentration of the lysates was determined by the Bradford method using Pierce(tm) BCA Protein Assay Kit (Thermo Scientific, USA).

### Isolation of CFZ-binding proteins

The columns were equilibrated with PBS buffer pH 7.0. The CFZ-linked column and control column were loaded with the *M. tuberculosis* lysates (∼ 600 mg protein for each column) and incubated at room temperature for 2 hours. Then the columns were washed with PBS buffer to reduce nonspecific binding of proteins until the baseline was stable. The columns were eluted with PBS buffer containing 2 M NaCl. The eluted fractions were run on 4 - 20% SDS-PAGE gradient gels, followed by staining with Coomassie Blue.

### Identification of CFZ binding proteins from *M. tuberculosis*

The proteins eluted from the CFZ-agrose column and separated on the SDS-PAGE gel were excised and subjected to in-gel digestion with trypsin followed by analysis by MALDI-TOF, and in parallel, another gel containing the protein bands was transferred to PVDF membrane for N-terminal sequence to identify N-terminal sequencing was performed by Edman degradation method.

### Cloning, sequencing, expression and purification of *M. tuberculosis* integration host factor (mIHF, Rv1388)

The *mIHF,* encoding the *M. tuberculosis* integration host factor, was amplified by PCR from *M. tuberculosis* H37Rv genomic DNA using forward primer 5’-GCGCCCATGGCCCTTCCCCAGTTGACCG - 3’ containing a NcoI restriction site (underlined) and reverse primer 5’-GCATCTCGAGGGCGGAGCCGAACTTTTCCAGC - 3’ a XhoI restriction site (underlined). The PCR fragments were digested with NcoI and XhoI (New England BioLabs, Inc., Ipswich, MA, USA), and ligated to plasmid pET28a digested with the same enzymes to construct recombinant plasmid which was confirmed by DNA sequencing. The mycobacterial mIHF protein was overexpressed in log phase *E. coli* strain BL21 (DE3) after induction with IPTG (1 mM) at 37 °C for 3.5 hours. The supernatant containing the recombinant mIHF was purified on Ni-NTA agarose column. Immobilized recombinant mIHF protein was washed by a 20 - 50 mM imidazole gradient and eluted with buffer (20 mM Tris-HCl, 300 mM NaCl, pH 8.0 and 250 mM imidazole). The purified protein was dialyzed against 10 mM Tris-HCl buffer (pH 7.5) to remove imidazole and stored at −70 °C for further experiments.

### CFZ and mIHF interaction by gel shift assay

Binding reactions (10 μl) contained recombinant *M. tuberculosis* H37Rv mIHF 0.18 μg and different concentration of CFZ or control drug were incubated at 10 mM Tris-HCl, 10 mM MgCl_2_, 1 mM DTT, pH 7.9 at room temperature for 30 min. 2□μl of 5 × loading dye (50% glycerol, 0.25 M acetate-KOH pH 6.8 and trace amount methyl green) were added, tubes were centrifuged for 5□min at 5000□rpm and mIHF–CFZ complexes were separated in acidic native gels, which contained 0.375 M acetate-KOH, pH 4.3, 15% polyacrylamide and 10% glycerol for separating gel and 0.0625 M acetate-KOH pH 6.8, 3% polyacrylamide and 10% glycerol for stacking gel, in running buffer with 0.35 M β-alanine, 0.14 M acetic acid (pH 4.3). Samples were electrophoresed in the stacking gel at 5 mA and then run overnight at 3 mA per gel at room temperature, followed by silver staining. The gel shift results were repeated 3 times.

### mIHF and DNA interaction

mIHF and DNA interaction was analyzed with supercoiled or linearized form of pUC19 plasmid DNA (linearized with HindIII) and mIHF concentrations ranging from 3.125 ng to 200 ng incubated in a buffer containing 10 mM Tris-HCl, pH 7.9; 50 mM NaCl; 10 mM MgCl_2_; 1 mM DDT on ice for 15 min. The mIHF-DNA complex was analyzed on 0.5% agarose gel following by Ethidium Bromide (EtBr) staining.

### Inhibition of mIHf and DNA interaction by CFZ

The CFZ inhibition assay was investigated with different concentrations of CFZ (0.05, 25, 50, 100, and 800 μg/ml) in the above mIHF (50 ng/reaction) and DNA (50 ng/reaction) binding reaction. CFZ and recombinant H37Rv mIHF were incubated on ice for 30 min. Then mIHF alone or mIHF-CFZ complex was incubated with DNA on ice for 15 min. The mIHF-(CFZ)-DNA complex was analyzed on 0.5% agarose gel, running in 1× TBE buffer and followed by Ethidium Bromide staining.

### Localization of mIHF and gene expression interference by CFZ

To fuse red fluorescent protein (RFP) to the C terminal of mIHF, mIHF was PCR-amplified using *M. tuberculosis* H37Rv genomic DNA as the template and primer pairs as Rv1388F: 5’-GCGCTTAATTAAGAAGGAGATATACATATGGCCCTTCCCCAGTTGACCG-3’ a d Rv1388R: 5’-GATCAAGCTTGGCGGAGCCGAACTTTTCCAGCAGGG-3’. Red fluorescent protein (RFP) was amplified by PCR and primer pairs as RedF: 5’-CGGAAGCTTGCCTCCTCCGAGGACGTCATC-3’ and RedR: 5’-TTCGATATCTTATGCTGCTGCTGCTGCTGCCAGG-3’. Rv1388 and RFP fragments were digested with PacI/HindIII and HindIII/EcoRv, respectively, then cloned into the tetracycline-inducible plasmid pUV15tetORM^31^ digested with PacI and EcoRV, resulting in the plasmid pUV15-mIHF-RFP. Then, recombinant constructs were verified by DNA sequencing and transformed into *M. tuberculosis* H37Ra as described^32^. Actively growing *M. tuberculosis* H37Ra/pUV15tet-mIHF-RFP was exposed with different concentration CFZ at 37°C for 72 hours and then ATc was added to induce the fusion protein expression. The *M. tuberculosis* cells expressing the pUV15tet-mIHF-RFP constructs were visualized by using a Nikon Eclipse E800 fluorescence microscope with a SPOT slider color camera at different time points.

## Acknowledgements

The work was supported in part by NIH grants AI099512 and AI108535.

## References

1 Barry, V. C. et al. A new series of phenazines (rimino-compounds) with high antituberculosis activity. Nature 179, 1013–1015 (1957).

2 Clofazimine (Lamprene) in leprosy. Drug Ther Bull 8, 7–8 (1970).

3 WHO. WHO treatment guidelines for drug-resistant tuberculosis, 2016 update. (2016).

4 Tam, C. M., Yew, W. W. & Yuen, K. Y. Treatment of multidrug-resistant and extensively drug-resistant tuberculosis: current status and future prospects. Expert Rev Clin Pharmacol 2, 405–421, doi:10.1586/ecp.09.19 (2009).

5 Cholo, M. C., Steel, H. C., Fourie, P. B., Germishuizen, W. A. & Anderson, R. Clofazimine: current status and future prospects. J Antimicrob Chemother 67, 290- 298, doi:10.1093/jac/dkr444 (2012).

6 O'Donnell, M. R., Padayatchi, N. & Metcalfe, J. Z. Elucidating the role of clofazimine for the treatment of tuberculosis. Int J Tuberc Lung Dis 20, 52–57, doi:10.5588/ijtld.16.0073 (2016).

7 Morrison, N. E. & Marley, G. M. Clofazimine binding studies with deoxyribonucleic acid. Int J Lepr Other Mycobact Dis 44, 475–481 (1976).

8 Barry, V., Belton, JG, Conalty, ML, et al.,. A New Series of Phenazines (Rimino-Compounds) With High Antituberculosis Activity. Nature 179, 1013–1015 (1957).

9 Van Rensburg, C. E., Joone, G. K., O'Sullivan, J. F. & Anderson, R. Antimicrobial activities of clofazimine and B669 are mediated by lysophospholipids. Antimicrob Agents Chemother 36, 2729–2735 (1992).

10 Yano, T. et al. Reduction of clofazimine by mycobacterial type 2 NADH:quinone oxidoreductase: a pathway for the generation of bactericidal levels of reactive oxygen species. J Biol Chem 286, 10276–10287, doi:10.1074/jbc.M110.200501 (2011).

11 Shi, W. et al. Pyrazinamide inhibits trans-translation in Mycobacterium tuberculosis. Science 333, 1630–1632, doi:10.1126/science.1208813 (2011).

12 Cole, S. T. et al. Deciphering the biology of Mycobacterium tuberculosis from the complete genome sequence. Nature 393, 537–544, doi:10.1038/31159 (1998).

13 Mishra, A. et al. Integration host factor of Mycobacterium tuberculosis, mIHF, compacts DNA by a bending mechanism. PloS one 8, e69985, doi:10.1371/journal.pone.0069985 (2013).

14 Sharadamma, N. et al. Molecular dissection of Mycobacterium tuberculosis integration host factor reveals novel insights into the mode of DNA binding and nucleoid compaction. The Journal of biological chemistry 289, 34325–34340, doi:10.1074/jbc.M114.608596 (2014).

15 Pedulla, M. L., Lee, M. H., Lever, D. C. & Hatfull, G. F. A novel host factor for integration of mycobacteriophage L5. Proceedings of the National Academy of Sciences of the United States of America 93, 15411–15416 (1996).

16 Reisfeld, R. A., Lewis, U. J. & Williams, D. E. Disk electrophoresis of basic proteins and peptides on polyacrylamide gels. Nature 195, 281–283 (1962).

17 David, H. L., Rastogi, N., Clavel-Seres, S. & Clement, F. Studies on clofazimine-resistance in mycobacteria: is the inability to isolate drug-resistance mutants related to its mode of action? Zentralblatt fur Bakteriologie, Mikrobiologie, und Hygiene. Series A, Medical microbiology, infectious diseases, virology, parasitology 266, 292–304 (1987).

18 Morrison, N. E. & Marley, G. M. The mode of action of clofazimine DNA binding studies. Int J Lepr Other Mycobact Dis 44, 133–134 (1976).

19 Ammerman, N. C. et al. Clofazimine has delayed antimicrobial activity against Mycobacterium tuberculosis both in vitro and in vivo. J Antimicrob Chemother 72, 455–461, doi:10.1093/jac/dkw417 (2017).

20 N. Odermatt, M. L., T. Herrmann, N. Dhar, S. Cole. Characterization of the Mycobacterial Integration Host Factor. EMBO Conference:“Tuberculosis 2016: Interdisciplinary research on tuberculosis and pathogenic mycobacteria (Pairs, France). Page 107 (2016).

21 Makafe, G. G. et al. Role of the Cys154Arg Substitution in Ribosomal Protein L3 in Oxazolidinone Resistance in Mycobacterium tuberculosis. Antimicrobial agents and chemotherapy 60, 3202–3206, doi:10.1128/AAC.00152-16 (2016).

22 Palmer, A. C. & Kishony, R. Opposing effects of target overexpression reveal drug mechanisms. Nat Commun 5, 4296, doi:10.1038/ncomms5296 (2014).

23 Zhang, S. et al. Identification of novel mutations associated with clofazimine resistance in Mycobacterium tuberculosis. J Antimicrob Chemother 70, 2507–2510, doi:10.1093/jac/dkv150 (2015).

24 Hartkoorn, R. C., Uplekar, S. & Cole, S. T. Cross-resistance between clofazimine and bedaquiline through upregulation of MmpL5 in Mycobacterium tuberculosis. Antimicrob Agents Chemother 58, 2979–2981, doi:10.1128/AAC.00037-14 (2014).

25 Almeida, D. et al. Mutations in pepQ Confer Low-level Resistance to Bedaquiline and Clofazimine in Mycobacterium tuberculosis. Antimicrob Agents Chemother, doi:10.1128/AAC.00753-16 (2016).

26 Pedulla, M. L. & Hatfull, G. F. Characterization of the mIHF gene of Mycobacterium smegmatis. J Bacteriol 180, 5473–5477 (1998).

27 Sassetti, C. M., Boyd, D. H. & Rubin, E. J. Genes required for mycobacterial growth defined by high density mutagenesis. Mol Microbiol 48, 77–84 (2003).

28 Lin, J., Chen, H., Droge, P. & Yan, J. Physical organization of DNA by multiple non-specific DNA-binding modes of integration host factor (IHF). PloS one 7, e49885, doi:10.1371/journal.pone.0049885 (2012).

29 Le, S. et al. Mechanosensing of DNA bending in a single specific protein-DNA complex. Sci Rep 3, 3508, doi:10.1038/srep03508 (2013).

30 Zhang, S., Shi, W., Feng, J., Zhang, W. & Zhang, Y. Varying effects of common tuberculosis drugs on enhancing clofazimine activity in vitro. Emerg Microbes Infect 6, e28, doi:10.1038/emi.2017.24 (2017).

31 Ehrt, S. et al. Controlling gene expression in mycobacteria with anhydrotetracycline and Tet repressor. Nucleic Acids Res 33, e21, doi:10.1093/nar/gni013 (2005).

32 Shi, W. et al. Aspartate decarboxylase (PanD) as a new target of pyrazinamide in Mycobacterium tuberculosis. Emerg Microbes Infect 3, e58, doi:10.1038/emi.2014.61 (2014).

